# My host’s enemy is my enemy: plasmids carrying CRISPR-Cas as a defence against phages

**DOI:** 10.1101/2023.11.01.565096

**Authors:** Berit Siedentop, Dario Rüegg, Sebastian Bonhoeffer, Hélène Chabas

## Abstract

Bacteria are infected by mobile genetic elements like plasmids and virulent phages, and those infections significantly impact bacterial ecology and evolution. Recent discoveries reveal that some plasmids carry anti-phage immune systems like CRISPR-Cas, suggesting that plasmids may participate in the coevolutionary arms-race between virulent phages and bacteria. Intuitively, this seems reasonable as virulent phages kill the plasmid’s obligate host. However, the efficiency of CRISPR-Cas systems carried by plasmids can be expected to be lower than those carried by the chromosome due to continuous segregation loss, creating susceptible cells for phage amplification. To evaluate the anti-phage protection efficiency of CRISPR-Cas on plasmids, we develop a stochastic model describing the dynamics of a virulent phage infection against which a conjugative plasmid defends using CRISPR-Cas. We show that CRISPR-Cas on plasmids provides robust protection, except in limited parameter-sets. In these cases, high segregation favours phage outbreaks by generating a population of defenceless cells on which the phage can evolve and escape CRISPR-Cas immunity. We show that the phage’s ability to exploit segregation loss depends strongly on the evolvability of both CRISPR-Cas and the phage itself.

## 1 Introduction

Bacteria are frequently infected by mobile genetic elements such as conjugative plasmids and virulent phages. These two types of mobile genetic elements spread in bacterial populations, and they are both dependent on their bacterial host to complete their life cycle. For the host, the fitness consequences of an infection depend on the type of mobile genetic element. On one hand, conjugative plasmid infection usually decreases the host’s fitness [1], but plasmids can encode accessory genes, such as antibiotic resistance, that can provide the host a fitness advantage in certain environments. On the other hand, successful virulent phage infections always result in cell death [2].

The evolutionary ecology between bacteria and phages or plasmids has been extensively studied. On the phage side, it is known that an important part of their coevolution with bacteria is mediated by immune systems, such as CRISPR-Cas, which are set of genes whose main function is to detect a phage infection and to protect the population against it [3]. Plasmids also interact with bacterial immune systems, but generally less is known about the evolutionary consequences of those immune systems for plasmids. However, it is known that plasmid maintenance strongly depends on genes responsible for replication and conjugation, as well as on accessory genes. In greater detail, plasmid maintenance depends on 1) the cost/benefit it brings to its host, 2) its rate of spreading horizontally through a mechanism called conjugation and 3) its ability to spread vertically during cell division. Importantly, this vertical transmission is imperfect, and a small fraction of daughter cells will not inherit the plasmid (segregation loss) [4].

Yet, much less is known about the interactions between plasmids and virulent phages. Virulent phages and conjugative plasmids frequently interact in natural environments as they are both highly prevalent in bacterial populations [5]. Intuitively, one can expect virulent phage-plasmid interactions to be amensal: even if the phage’s fitness is not modified by the presence of the plasmid, a virulent phage infection results in a cost for the plasmid, as it cannot survive the death of its host. There is evidence that under certain conditions a virulent phage infection can lead to the exclusion of an established, costly plasmid, exemplifying the evolutionary pressure phages inflict on plasmids [6]. Therefore, one can expect that plasmids would benefit of evolving ways of protecting themselves (and consequently their host) against virulent phages.

Indeed, there is evidence that some plasmids carry systems to defend against phages. For example, plasmids frequently carry toxin-antitoxin systems, and some of these toxin-antitoxin systems have been demonstrated to participate in phage inhibition [7–10]. Recent bioinformatic studies show that toxin-antitoxin systems are not the only phage defence mechanism carried by plasmids: a small, but still significant proportion of plasmids carry CRISPR-Cas systems [11]. CRISPR-Cas systems are a prevalent and well-known form of adaptive immunity that bacteria use to defend against phages [3]. They consist of two loci, a Cas locus that codes for all the proteins the system needs to function, and a CRISPR locus, which can be seen as a library of small sequences (called spacers) acquired from previous invaders (such as phages or plasmids). Cas proteins use these spacers as guides to trigger the degradation of the foreign genome they target. About a third of the spacers carried by CRISPR-Cas on plasmids are predicted to target phages [11], raising the hypothesis that these systems can be used by plasmids to defend against phages.

However, there are reasons to hypothesize that CRISPR-Cas systems on plasmids may be unable to provide as efficient immunity against virulent phages as chromosomal CRISPR-Cas, because plasmids experience segregation loss. Indeed, immune loss is known to favour coexistence between phages and bacteria carrying CRISPR-Cas by increasing phage reproduction [12]. Additionally, because a single point mutation in the phage protospacer (the phage sequence targeted by the spacer) is often enough to escape a spacer [13], segregation loss could result in a higher probability of escape, as the phage could evolve on this susceptible population [14]. Nevertheless, how relevant segregation loss is in the complex tripartite interaction between CRISPR-Cas, plasmids and phages is unclear.

To test whether CRISPR-Cas carried by plasmids can provide the same anti-phage protection efficiency as chromosomal CRISPR-Cas and to identify key factors in this complex tripartite interaction that influence phage protection, we developed a stochastic model describing phage, bacterial, plasmid and CRISPR-Cas dynamics. We show that in most biologically relevant parameter sets, CRISPR-Cas on plasmids provides as efficient immunity as chromosomal CRISPR-Cas, further strengthening the evidence that plasmids can contribute to phage immunity of bacterial cells. In rare cases, CRISPR-Cas on plasmids provide a less efficient immunity than chromosomal CRISPR-Cas. We demonstrate that this reduced efficiency is due to segregation loss, which produces a subpopulation of susceptible cells that the phage can exploit to facilitate the escape of CRISPR-Cas immunity. Yet, whether segregation loss leads to a decreased CRISPR-Cas efficiency is strongly dependent on the phage’s evolvability and the ability of CRISPR-Cas to acquire new spacers.

## 2 Model of plasmid, CRISPR-Cas and phage dynamics

Our mathematical model describes the potential of CRISPR-Cas systems carried by plasmids to protect a bacterial population from a phage outbreak (figure 1) in the early stages of an outbreak. Experimental evidence shows that in the first days of an outbreak, cells only acquire a single spacer against the phage, which is often sufficient to drive the phage to extinction [15]. Therefore, we assume in our model that each bacterial cell can only acquire a single spacer and each phage can only escape one spacer. If the phage is able to survive the initial phase of an outbreak, bacterial cells may acquire multiple spacers [16, 17]. However, our model does not consider this long-term dynamic. As a result, if multiple spacers per cell are necessary to drive the phage to extinction, our model predicts phage survival.

**Figure 1.**
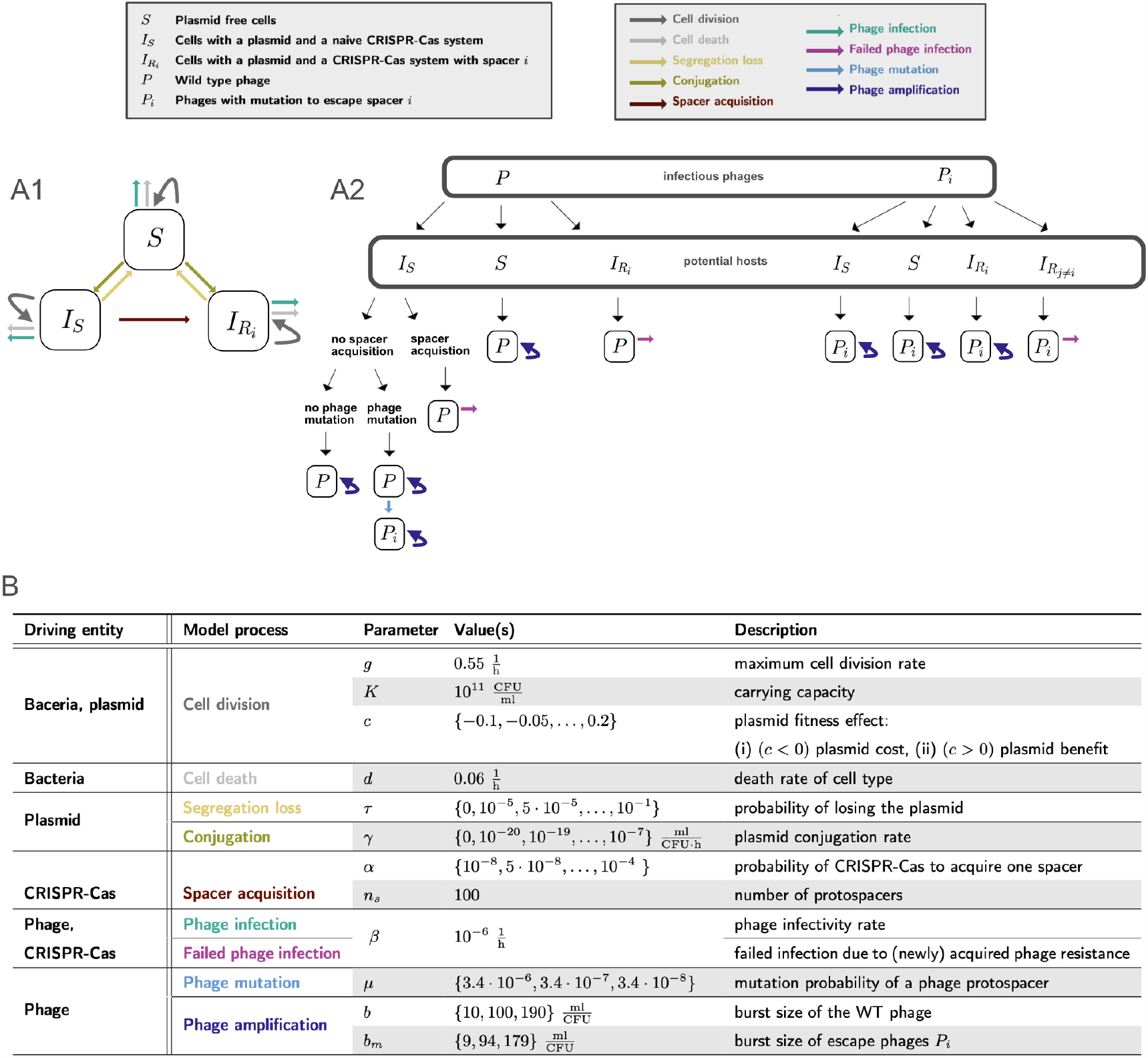
Visualisation of the modelled CRISPR-Cas, plasmid and phage dynamics and the corresponding parameters. **A1**. Schematic diagram of plasmid and CRISPR-Cas dynamics illustrating the processes described by equations 1-3. **A2**. Schematic diagram of phage infection and CRISPR-Cas processes described by equations 1-5. **B**. Overview of the model processes and associated parameters including used parameter values. The choice of model parameters is specified in the main text in section 2.5.

Our model is based on the model for type I/II CRISPR-Cas systems of Chabas et al., 2022 [18], with the addition of basic plasmid processes incorporated along the lines of Igler et al., 2022 [19]. We chose to limit our analysis on type I CRISPR-Cas systems because type I CRISPR-Cas systems are the most frequent on bacterial chromosomes and plasmids [11, 20]. Our model considers that a phage contains multiple protospacers and that CRISPR-Cas will randomly pick between them each time a spacer is acquired [21, 22]. This results in a population-wide spacer diversity, which has been previously shown to be critical for the efficiency of CRISPR-Cas against evolving phages [14, 15, 18]. In our model, we assume that the maximal number of different protospacers - and therewith spacers within the bacterial population - is limited by *n*_*S*_ = 100. This number of spacers is sufficiently high to lead to phage to extinction [18], allowing us to study the various potential outcomes of a phage outbreak.

We consider one type of plasmid with a CRISPR-Cas system, where plasmid copies can only differ in the CRISPR-Cas spacer content. As we are interested in phage protection, we only explicitly trace phage-specific spacers in our model. We assume that each CRISPR-Cas system is initially naive to the phage and can only acquire one spacer, as we only model the initial dynamics of a phage outbreak [18]. Therefore, the bacterial population consists of the following compartments: (i) *S*: plasmid free cells, (ii) *I*_*S*_: bacterial cells infected with a plasmid carrying CRISPR-Cas system, that is naive to the phage and (iii) 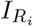: cells infected with a plasmid carrying a CRISPR-Cas system with a phage-specific spacer *i*.

### 2.1 Bacterial population size

Bacterial division is modelled by a density-dependent division rate 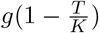, where *g* is the maximal division rate, 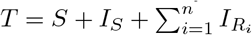 the total cell density and *K* the carrying capacity. The plasmid is associated with a fitness effect *c* (*c >* 0 representing a cost and *c <* 0 a benefit), which modifies the maximum division rate of plasmid carrying cells by the factor (1 − *c*). Furthermore, we assume that bacteria die at a constant rate *d*. Altogether we obtain a model with a bounded bacterial population size, where the plasmid’s fitness effect remains effective even after this bound is reached. The plasmid fitness effect influences only the rate of division but not the bounded bacterial population size.

### 2.2 Segregation loss and conjugation

During cell division, plasmid carrying cells can lose the plasmid due to segregation loss with a probability *τ*. We assume a well-mixed population where plasmid conjugation depends on host cell density and occurs at a rate *γ*. As plasmids commonly carry exclusions systems [23], which prevent the conjugation of a plasmid into cells infected by the same plasmid variant, we exclude plasmid co-infection.

### 2.3 Phage infection and spacer acquisition

If a wild type (WT) phage *P* infects a cell *S* which does not carry a plasmid and therefore has no CRISPR-Cas system, the phage can successfully infect with rate *β* and amplify. During this amplification, each of the *n*_*S*_ protospacers of the WT phage can mutate with probability *μ*. We assume that a phage can only have one mutated protospacer and that mutations are not reversible. An infection of *S* by the WT phage leads to either *b* number of progeny WT phages, or in case of mutation, both the escape phage *P*_*i*_ and the WT phage *P* amplify. We assume in this case, as Chabas et al., 2022 [18], that *P* and *P*_*i*_ each produce half of their respective number of progeny: 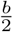 and 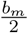. Additionally, we assume that the escape phages have lower fitness than the WT phage, i.e. *b*_*m*_ *< b* as demonstrated experimentally for one model system [18, 24]. In case *P* infects a cell carrying a plasmid with a naive CRISPR-Cas system *I*_*S*_ the phage can either successfully infect and amplify (with opportunity for phage mutation), or the naive CRISPR-Cas system can acquire a spacer with a probability *α*, the then resistant cell survives the infection and no phage gets amplified. If a WT phage infects 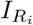, a cell which already carries spacer *i*, the phage infection fails. A failed phage infection leads to degradation of the phage without any consequences for the bacterial cell.

Escape phages *P*_*i*_ can successfully infect plasmid-free cells *S* and cells with a naive CRISPR-Cas system *I*_*S*_, which leads to cell death and *b*_*m*_ number of progeny. Additionally, escape phages *P*_*i*_ (with an escape mutation in protospacer *i*) can also successfully infect cells 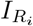, which carry spacer *i*. However, if the escape phage has an irrelevant mutation in protospacer *j* for the infection of 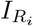, i.e. *j* ≠ *i*, the phage infection fails.

### 2.4 Further simplifications to model the biological processes

For simplicity, we do not consider spacer acquisition of mutated phage sequences, but only of WT sequences. This event would only occur when *P*_*i*_ infects *I*_*S*_, with a probability 1*/n*_*s*_ in case of spacer acquisition. As in our model *n*_*s*_ equals 100, such an event is rare.

Furthermore, we disregard the possibility of within cell spacer heterogeneity. Such heterogeneity could stem from multicopy plasmids, where each copy of a plasmid carries a CRISPR-Cas system. In our model we do not track plasmid copy number explicitly. In theory, each copy of the CRISPR-Cas system could acquire distinct spacers during a single phage infection, leading to within cell spacer heterogeneity. However, if we assume that for each CRISPR-Cas system the probability of spacer acquisition is equal to *α* ≤ 10^*−*4^ and assume independent acquisition for each CRISPR-locus, this event is extremely rare. Additionally, we exclude the possibility of within spacer heterogeneity acquired through conjugation, as surface exclusion systems are believed to be common [25], which prevent a plasmid of the same exclusion class to enter the cell twice. Spacer heterogeneity could also stem from phage reinfection of a cell that already contains a spacer. However, for chromosomal CRISPR-Cas systems, there is some experimental evidence showing that cells with multiple spacers are rare during the initial phase of a phage outbreak.

We also do not consider natural phage decay as our model is designed to study the initial dynamics between CRISPR-Cas and a virulent phage. At that time scale, phage natural decay is negligible under laboratory conditions.

Besides capturing the dynamics of CRISPR-Cas on plasmids, our mathematical model also describes chromosomal CRISPR-Cas when all plasmid parameters are set to zero, i.e. *γ* = *c* = *τ* = 0.

The above described processes can be formalised with the following differential equations:

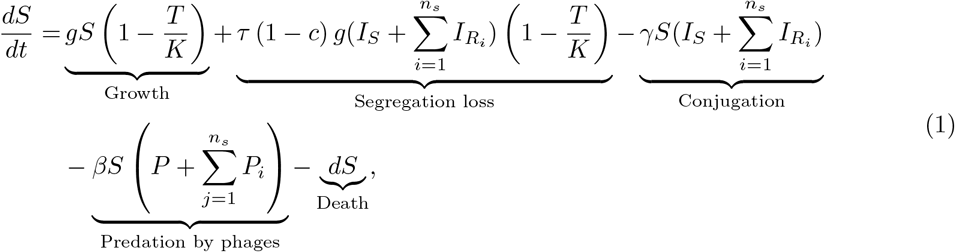

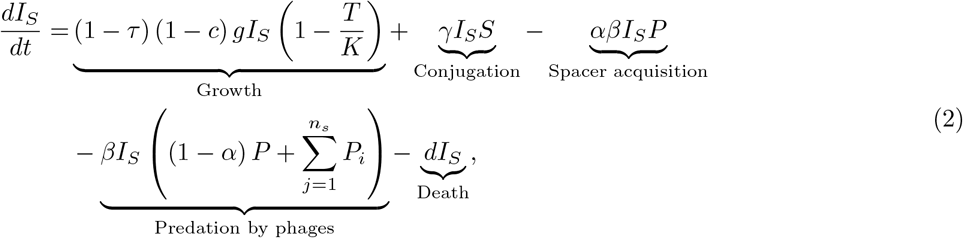

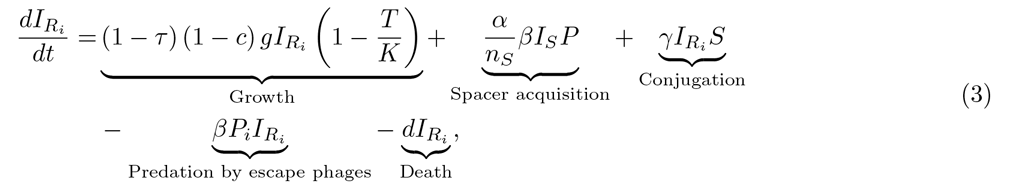

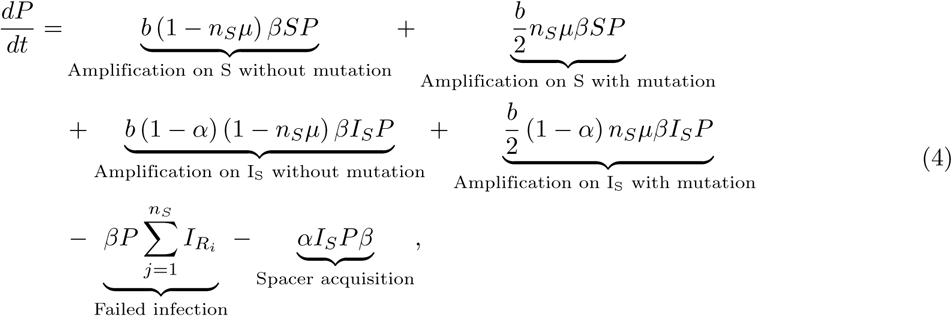

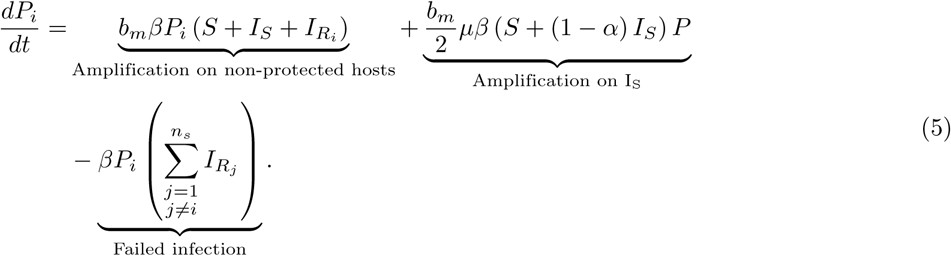

### 2.5 Implementation

For our investigations, we simulated the equations 1-5 stochastically. We terminated our simulations if either the phage or bacterial population went extinct. Although we had a maximum time limit of 150 hours to track the populations, it was never reached. For the implementation we used the *adaptivetau* package (version 2.2-3) in R (version 3.6.0), which implements the Gillespie algorithm with adaptive *τ* -leaping [26, 27]. For comparison, we ran simulations where the CRISPR-Cas system was located on a plasmid as well where it was located on the chromosome. For each parameter set, given by the parameter combinations in 1B (with fixed pairings of phage burst sizes: (*b, b*_*m*_) = {10, 9}, {100, 94} or {190, 179}), we simulated 100 runs, only for cases where the CRISPR-Cas system was located on the chromosome (*γ, τ, c* = 0) we ran 10000 simulations. For chromosomal CRISPR-Cas, we increased the number of simulations as we compared several parameter sets of CRISPR-Cas on plasmids, which differed in plasmid conjugation, segregation loss, and fitness effect, to one chromosomal CRISPR-Cas. All simulations were started with the following initial conditions: *I*_*S*0_ = 10^6^ and *P*_0_ = 10^5^, reflecting a phage infection of a growing bacterial population, where initially all bacteria carry a plasmid with a naive CRISPR-Cas system. CRISPR-Cas related parameters were chosen in line with Chabas et al., 2022 [18]. The choice of bacterial growth parameters was based on Chabas et al., 2022 [18], but differs slightly due to different ways of modelling bacterial growth and death. We chose the ranges of phage mutation rate and of phage escape cost based on experimental data from Chabas et al., 2019 [24] (we assume that the measured cost only results from a decreased burst size). The range of phage burst size accounts for phages with a high burst size (e.g. [28]) but also acknowledge that virulent phages can have a very small burst size on certain hosts (e.g. [29–31]). The parameter ranges for the plasmid conjugation rates and plasmid fitness effects were chosen in line with two meta-analysis on those two respective parameters [1, 32]. We used the same parameter range for segregation loss as Lethinen et al., 2021 [33]. For our simulations we explored the parameter range in equal linear or linear-log interval steps, with a particular focus on plasmid parameters.

### 2.6 Identification of key parameters determining efficiency of CRISPR-Cas on plasmids

To estimate the association between CRISPR-Cas efficiency and CRISPR-Cas location, we calculated the risk difference (RD) of CRISPR-Cas phage protection efficiency:

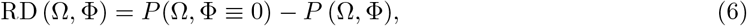

where *P* denotes the probability of phage extinction dependent on the chosen parameter set. Ω = {*α, μ, b, b*_*m*_} is the parameter set of CRISPR-Cas and phage characteristics and Φ = {*γ, c, τ*} the parameter set of plasmid characteristics used in the simulations (values given in figure 1B). Simulations with Φ ≡ 0 represent chromosomal simulations. Within a given parameter set (Ω, Φ), the probability of phage extinction was calculated by dividing the number of simulations where the phage went to extinction by the total number of simulations (plasmid: 100, chromosome: 10000). For each RD, we calculated the corresponding 95% confidence intervals. Based on the 95% confidence intervals, we assumed an association between the location of CRISPR-Cas and phage protection efficiency. The CRISPR-Cas performance on a plasmid was considered worse than on the chromosome if the lower bound of the 95% confidence interval was above zero, better in case the upper bound was below zero, and the same if the 95% confidence intervals included zero. To explore the relevance of CRISPR-Cas, plasmid and phage characteristics on the performance of CRISPR-Cas on plasmids, we used a supervised learning model based on boosted gradients trees. The implementation was done with the R package eXtreme Gradient Boost (XGBoost Verison 1.6.0.1 and R 4.2.1 version) [34], which relies on ensembles of trees. Further details about the concept of gradient boosted trees and algorithm details can be found in the XGBoost documentation [35]. We used this supervised learning technique based on trees to accurately capture non-linear effects. We fitted a statistical model with XGBoost predicting CRISPR-Cas performance of systems located on the plasmid based on the above described categories: better, same, worse. We used CRISPR-Cas, plasmid and phage parameters as predictors for the statistical model.

We aimed to restrict our investigation to biological relevant parameter ranges, as the assessment of parameter importance is dependent on the choice of the parameter space under investigation. The choice of parameter ranges is explained in section 2.5.

## 3 Results

### 3.1 Plasmids can efficiently protect against virulent phages using CRISPR-Cas

To determine whether plasmids can efficiently defend against virulent phages using their CRISPR-Cas systems, we implemented a stochastic model describing phage, plasmid and CRISPR-Cas dynamics. From this model, we extract the probability for a CRISPR-Cas system carried by a plasmid or by the chromosome to lead the phage to extinction (hereafter called CRISPR-Cas efficiency). We run this model using a range of plasmid, CRISPR-Cas and phage parameters that are biologically realistic (figure 1B). For each parameter set, we calculate the risk difference (RD) between the efficiency of chromosomal CRISPR-Cas and systems on plasmids (equation (6)) to quantify the association between the location of CRISPR-Cas and the protection efficiency. In total, we calculate the risk difference for 75528 different parameter sets and the 95% confidence intervals. We assume an association between CRISPR-Cas efficiency and CRISPR-Cas location when the 95% interval does not include the value zero. For 84% of the parameter sets, we find no association between CRISPR-Cas efficiency and CRISPR-Cas location, which we categorize as “same”. Only for 16% of our simulated parameter-sets we observe an association between location and CRISPR-Cas efficiency. In cases where we observe a difference in efficiency, CRISPR-Cas on plasmids is less efficient for 14% of the parameter sets, which we categorize as “worse”, and more efficient for 2% of the parameter sets, which we categorize as “better” (figure 2). Since the percentage of parameter sets categorized as “better” is very low (less than 2.5%), it is consistent with the hypothesis that those parameter sets do no differ significantly from those categorized as “same”. We conclude that for most of our investigated parameter sets, plasmids carrying CRISPR-Cas can defend as efficiently against phages as chromosomes carrying CRISPR-Cas.

**Figure 2.**
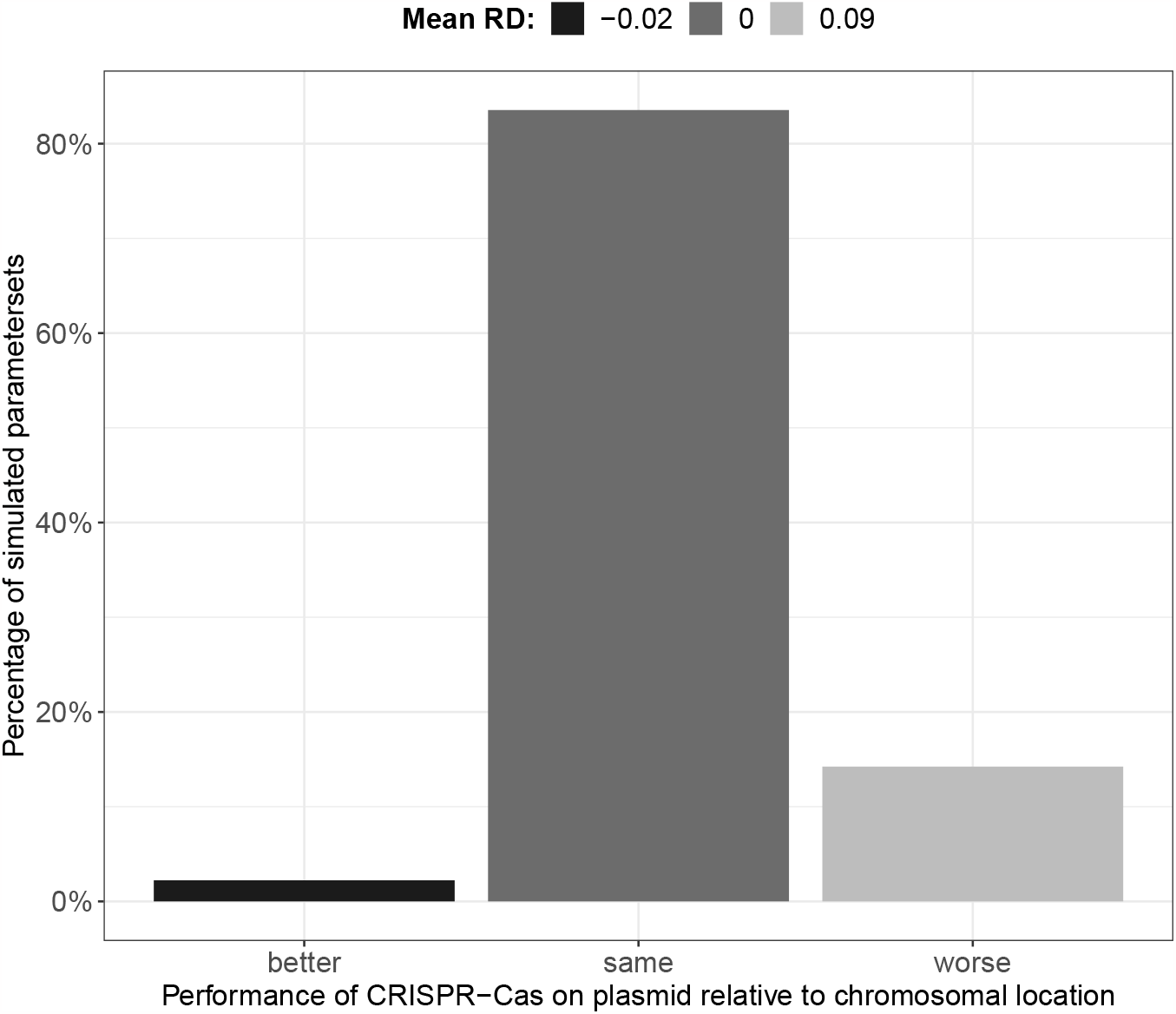
Anti-phage protection performance of plasmids using CRISPR-Cas. Percentage of all simulated plasmid parameter sets stratified according to CRISPR-Cas performance in comparison to chromosomal CRISPR-Cas. The mean RD between CRISPR-Cas efficiency on the chromosome and CRISPR-Cas efficiency on the plasmid is shown for each category.

To gain better insight into the differences in CRISPR-Cas efficiency, we evaluate by how much the protection differs when plasmids provide altered protection efficiency. To do so, we calculate the mean risk difference (RD) for each group of parameter sets categorized as “worse”, “same” or “better”. For the parameter sets categorized as “worse”, we obtain a mean RD of 0.09, meaning that CRISPR-Cas on plasmids have an absolute decrease in efficiency of 9% compared to chromosomal CRISPR-Cas. The mean RD for parameter sets categorized as “same” is zero and the mean RD for parameter sets categorized as “better” is −0.02, meaning that CRISPR-Cas on plasmids have an absolute increase in efficiency of 2% compared to chromosomal CRISPR-Cas (figure 2). We conclude that the difference in efficiency between CRISPR-Cas on plasmids and chromosomal CRISPR-Cas is the biggest when the plasmid provides weakened protection.

So far, we show that in 16% of our simulations with CRISPR-Cas on plasmids, the efficiency in immunity conferred by CRISPR-Cas on plasmids differs from the one conferred by a chromosomal CRISPR-Cas. However, it is unclear if the phage, CRISPR-Cas, or the plasmid influences this difference in immune efficiency the most. To determine this, we train a statistical classification model based on boosted gradient trees (further details in supplement section 1). We observe that the model’s most important parameters to distinguish between parameter sets where the plasmid provides “worse”, “same” or “better” efficiency, are the probabilities of 1) spacer acquisition *α*, 2) segregation loss *τ*, and 3) phage’s protospacer mutation *μ* (figure 3A). We conclude that the outcome of this tripartite dynamic depends on all 3 entities, and that none can be neglected. How those parameters influence the ability of plasmids to defend against phages with CRISPR-Cas is investigated in the following.

**Figure 3.**
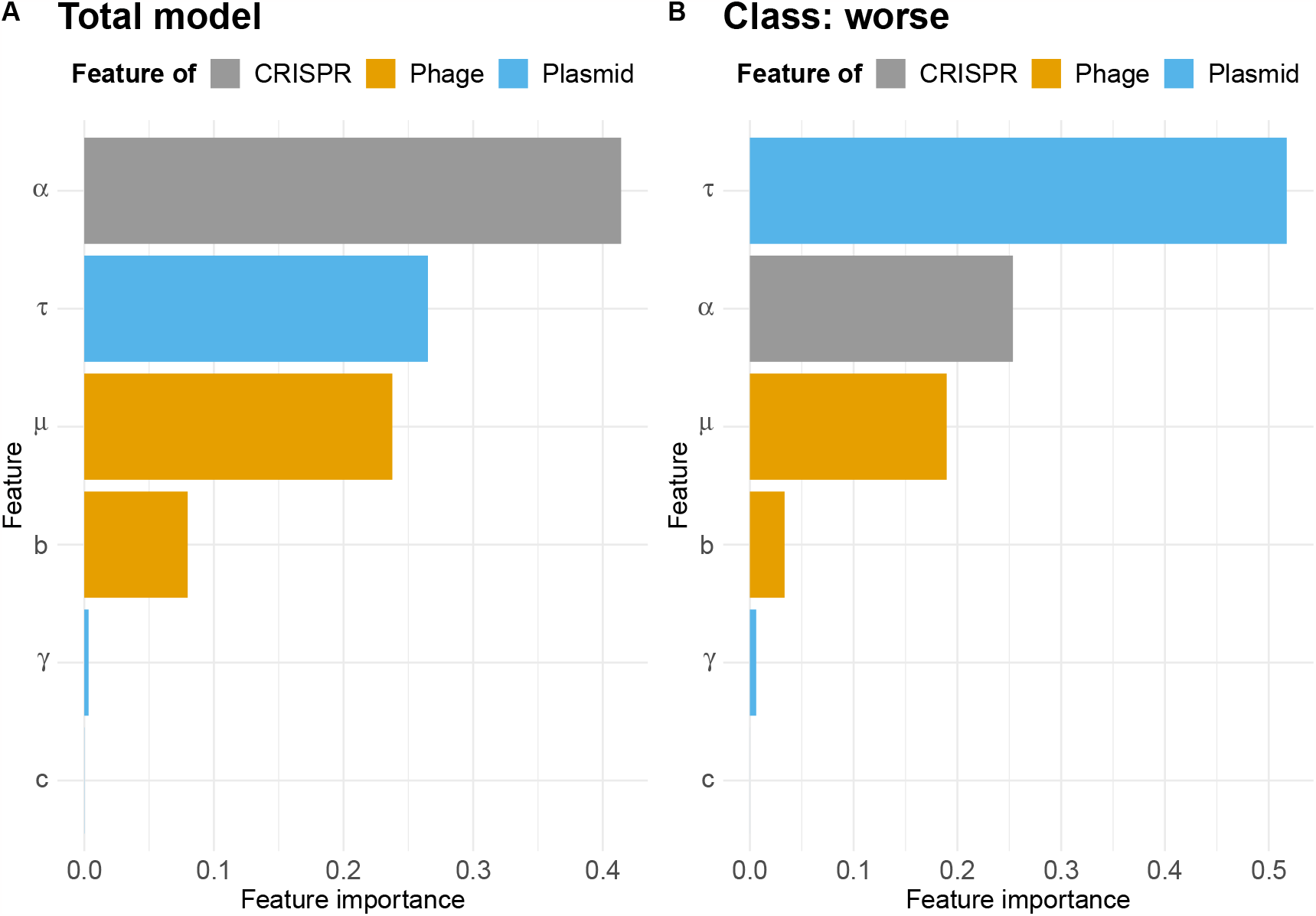
Importance of biological parameters to predict CRISPR-Cas performance on plasmids. **A**. Global importance of parameters in classifying the CRISPR-Cas efficiency on plasmids into the three categories “worse”, “same” and “better”. **B**. The model’s most important parameters in determining when the CRISPR-Cas efficiency on plasmids is worse than the efficiency of chromosomal systems.

### 3.2 Plasmid segregation loss favours phage survival by facilitating evolutionary emergence

So far, we show that the probabilities of i) spacer acquisition, ii) segregation loss and iii) phage protospacer mutation are most important to predict the efficiency of plasmids in defending against phages using CRISPR-Cas. Because immune loss is known to favour phage survival [12], we hypothesize that segregation loss could be the most important parameter to explain cases when CRISPR-Cas on plasmids are less efficient. To test for this, we determine which parameters are used by the previously described classification model to identify the “worse” class (Figure 3B), and indeed confirm that segregation loss is the most important parameter to determine when CRISPR-Cas on plasmids perform worse. Therefore, we now focus on investigating how segregation loss weakens the ability of plasmids to protect against phages using CRISPR-Cas. We reason that there are three non-mutually exclusive mechanisms, which can explain this.

First, segregation loss could decrease the generated population-based spacer diversity, which is known to be key for phage survival [14, 15] *(spacer diversity hypothesis)*. During segregation loss, one daughter cell does not inherit the plasmid and its CRISPR-Cas system. This results in a smaller number of cells carrying CRISPR-Cas, and consequently, in a smaller number of cells that have the opportunity to acquire a spacer. This could lead to a lower spacer diversity, which favours phage survival.

Second, segregation loss generates a population of susceptible cells, which could be large enough to produce more phage virions than can be killed by cells carrying a spacer, resulting in phage survival *(phage amplification hypothesis)*.

Third, segregation loss could facilitate evolutionary emergence, i.e. outbreaks caused by phage escape mutants *(phage evolution hypothesis)*. In this scenario, there are more opportunities for the WT phage to evolve escape mutations as it has more susceptible cells to infect. Additionally, there is a higher survival probability for the newly generated escape mutants, as their host population is increased. This could bias the arms race between phage diversity and spacer diversity in favour of the phage.

We first test the validity of the *spacer diversity hypothesis* by assessing if an increase in segregation loss decreases the initial spacer diversity (spacer diversity when no naive CRISPR-Cas system remains in the population), as after this time point, no new distinct spacers can be introduced in the population. We observe that the initial spacer diversity is not influenced by segregation loss (figure S1 C,D), ruling out the *spacer diversity hypothesis*.

To test the *phage amplification hypothesis*, we set the phage protospacer mutation probability to zero (*μ* = 0). By doing so, we test if the amplification of the wild type phage *P* without the evolution of escape mutants explains why segregation loss weakens the ability of plasmids to efficiently defend using CRISPR-Cas. In this case, we do not observe any decrease in phage protection efficiency while increasing segregation loss (figure 4B), which rules out the *phage amplification hypothesis*. This also shows that the susceptible population can only decrease CRISPR-Cas efficiency in case the phage has the ability to escape CRISPR-Cas.

**Figure 4.**
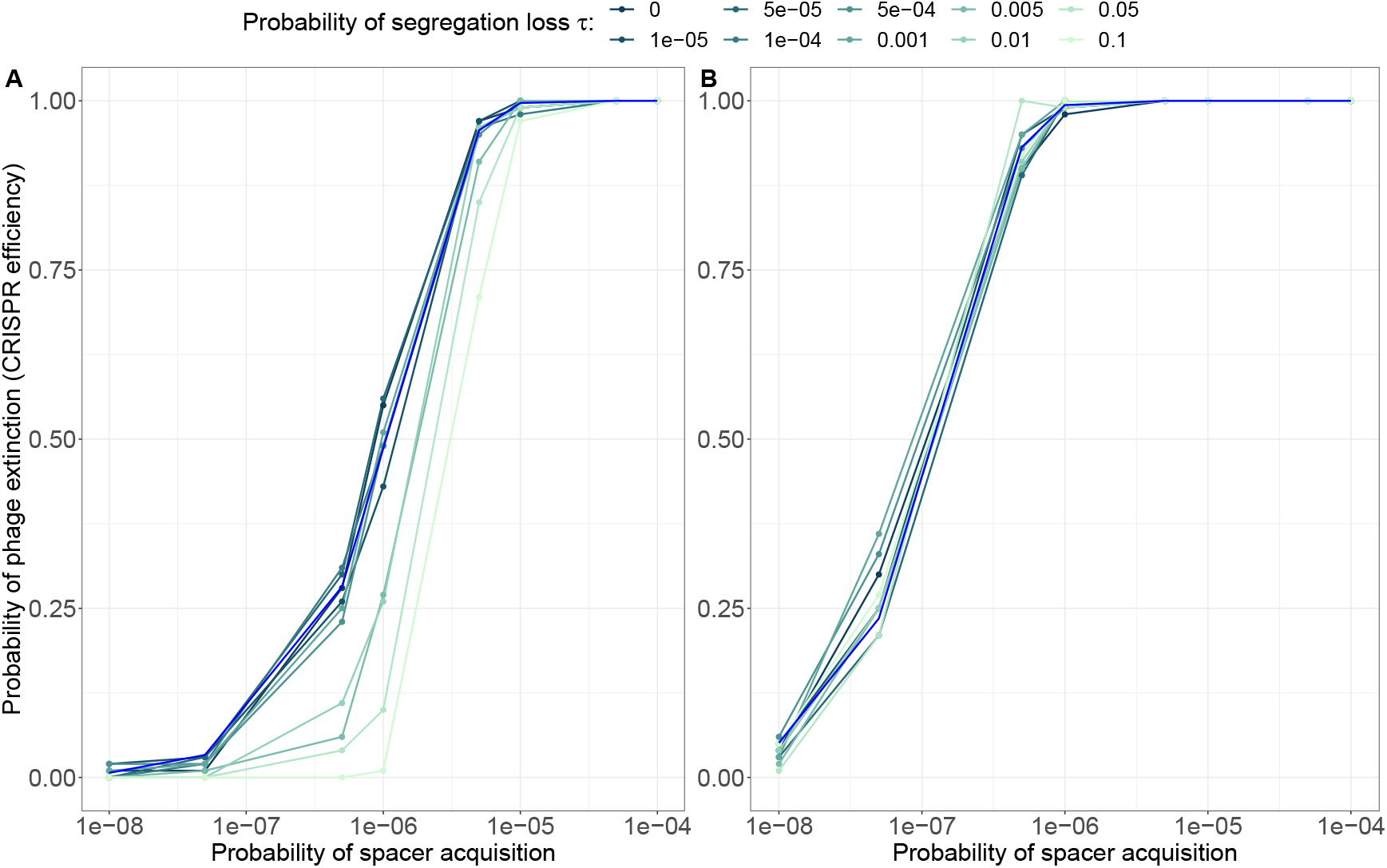
Impact of varying acquisition rates, segregation loss and phage mutation on the probability of phage extinction (CRISPR-Cas efficiency). **A** The phage has the possibility to escape acquired spacers by mutating protospacers with probability *μ* = 3.4 10^*−*7^. **B** The phage has no possibility to escape acquired CRISPR-Cas spacers *μ* = 0. The conjugation rate, plasmid cost and phage burs sizes are fixed at *γ* = 10^*−*14^, *c* = 0.05, *b* = 190 and *b*_*m*_ = 179. The probability of phage extinction for CRISPR-Cas systems on the chromosome is indicated in blue.

We show that segregation loss only leads to decreased CRISPR-Cas efficiency when the phage is able to escape CRISPR-Cas, which is consistent with the *phage evolution hypothesis*. To further investigate the *phage evolution hypothesis*, we analyze the proportion of escape phages both at the beginning and at the end of the phage outbreak, considering varying degrees of segregation loss. We observe that, during the very early stages of the outbreak, the proportion of escape mutants is consistently low, regardless of the level of segregation loss. However, at the end of the phage outbreak, we observe the dominance of escape mutants when segregation loss is high (figure S2). This confirms the *phage evolution hypothesis*, showing that a decreased CRISPR-Cas efficiency results from segregation loss favouring the outbreak of escape phages.

### 3.3 The benefit of segregation loss in facilitating evolutionary emergence is governed by CRISPR-Cas and phage characteristics

Overall, we show that plasmid, phage and CRISPR-Cas parameters are important to predict the performance of plasmids using CRISPR-Cas. So far, we only explain the role of segregation loss. We will not investigate the relevance of further plasmid parameters, i.e. conjugation and fitness effect, as our statistical model indicates that they only have minor relevance on CRISPR-Cas efficiency (figure 3). In the following we aim to understand the impact of the remaining CRISPR-Cas and phage parameters on the plasmid’s ability to defend against phages using CRISPR-Cas.

We first investigate the impact of CRISPR-Cas biology, which is represented in our model by the probability of spacer acquisition *α*, on the efficiency of CRISPR-Cas on plasmids. In line with previous studies, we show that the probability of spacer acquisition has a high impact on phage survival [18, 36, 37] and that efficiency differences between CRISPR-Cas on plasmids and chromosomal CRISPR-Cas occur especially at intermediate levels of spacer acquisition. Importantly, we find that even in cases where plasmids undergo high segregation loss, CRISPR-Cas on plasmid can provide the same phage protection as chromosomal CRISPR-Cas when spacer acquisition is high (figure 4A). In those cases the probability of acquiring a spacer (CRISPR-Cas evolutionary potential) is high enough to ensure evolutionary rescue of the bacterial population from the evolving phage population independent of the probability of segregation loss. At low acquisition probabilities, the phage is likely to kill the bacteria before CRISPR-Cas could acquire any spacer (figure S1A), making the presence of a sub-population for phage evolution redundant. Therefore, the presence of a sub-population of susceptible cell does not necessarily lead to an increase in phage survival.

Last, we explore the role of phage parameters, i.e. the protospacer mutation probability *μ* and the phage burst size *b*, on the efficiency of CRISPR-Cas on plasmids. We find that the phage’s evolutionary potential, i.e. its mutation probability and its burst size, can modify the benefit segregation loss provides to the phage. When the phage has a high protospacer mutation probability, this advantage in the co-evolutionary arms-race between CRISPR-Cas and phage can be sufficient to ensure phage survival and the phage is less dependent on segregation loss for its survival (figure 5A). However, in case the phage does not show high levels of mutations, segregation loss provides a benefit for phage survival and therefore decreases the efficiency of CRISPR-Cas on plasmids. Besides mutation, the phage burst size can also alter the evolutionary potential of phages. Indeed, we see in case the evolutionary potential is lowered by a small phage burst size, that segregation loss matters less for the efficiency of CRISPR-Cas on plasmids (figure 5B). In summary, we conclude that the evolutionary potentials of CRISPR-Cas and phage, which are determined in our model by the probabilities of acquiring a spacer or of mutating a protospacer and the phage burst size, can decrease the probability of evolutionary emergence, thus decreasing the impact of segregation loss on the immune efficiency a plasmid provides using CRISPR-Cas.

**Figure 5.**
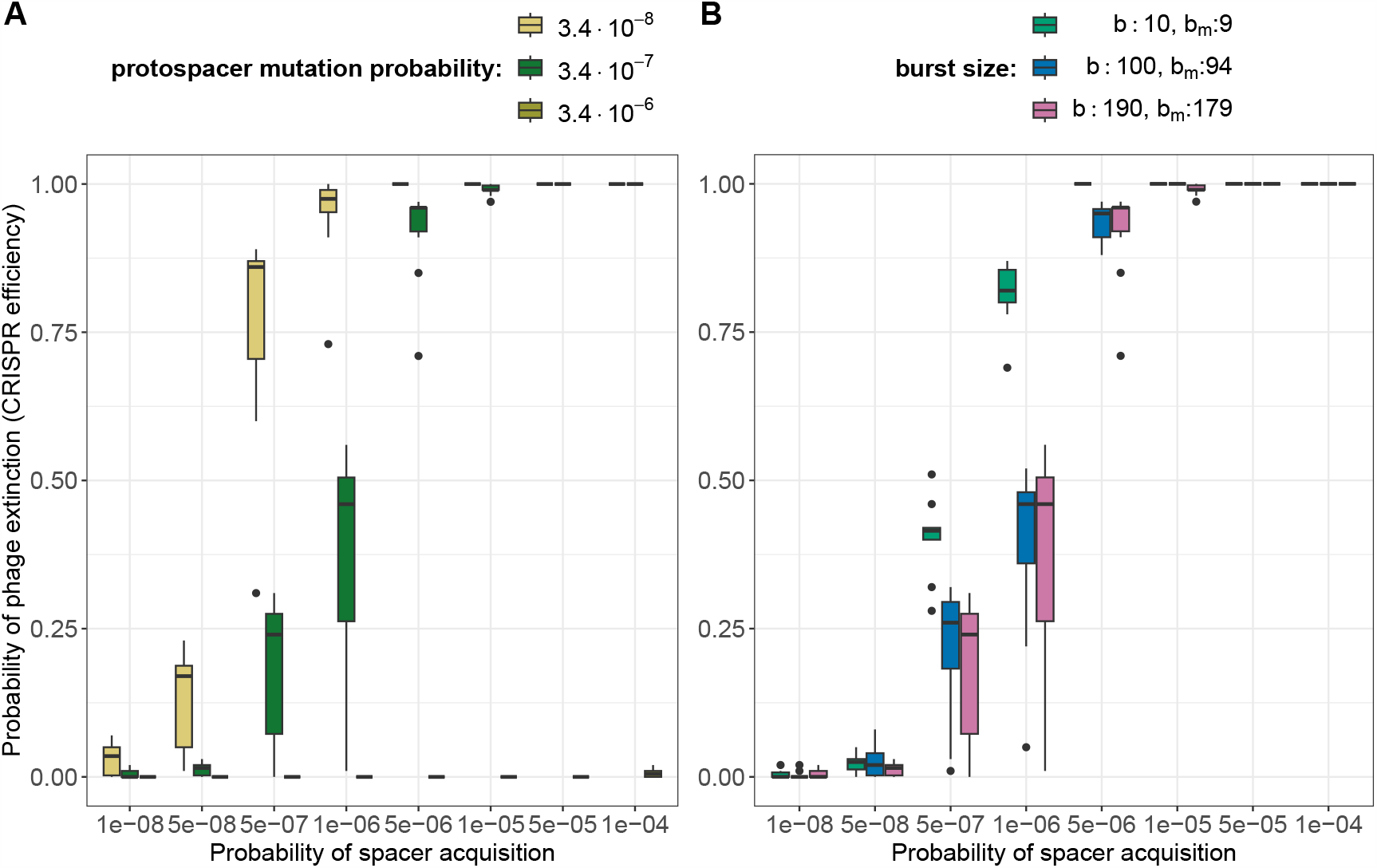
Impact of segregation loss on the probability of phage extinction (CRISPR-Cas efficiency) depending on phage characteristics. For different phage protospacer mutation probability (**A**) or phage burst sizes (**B**), we show the distribution of CRISPR-Cas efficiency across all the tested probabilities of segregation loss (see figure 1) for each probability of spacer acquisition. Hence, a wider boxplot indicates a greater impact of segregation loss on CRISPR-Cas efficiency. The conjugation rate and plasmid cost are fixed at *γ* = 10^*−*14^ and *c* = 0.05. The phage burst size is fixed in (**A**) to *b* = 190, *b*_*m*_ = 179 and the protospacer mutation probability in subplot (**B**) to *μ* = 3.4 *·* 10^*−*7^.

## 4 Discussion

In this work, we investigate if plasmids can provide an as efficient anti-phage immunity using CRISPR-Cas as chromosomal CRISPR-Cas systems. We show that, in 86% of the investigated parameter sets, CRISPR-Cas on plasmids provides at least the same protection efficiency as a chromosomal CRISPR-Cas system. However, for 14% of the tested parameter sets, we find a considerable decrease in efficiency for CRISPR-Cas on plasmids. Additionally, we show that in these limited sets of parameters, high segregation loss decreases CRISPR-Cas efficiency as it facilitates phage evolutionary emergence. However, high segregation loss does not necessarily lead to decreased efficiency of CRISPR-Cas on plasmids. Whether segregation loss alters the performance of CRISPR-Cas on plasmids is also strongly dependent on the phage and CRISPR-Cas. Overall, our results emphasize the importance of studying the interactions of multiple mobile genetic elements to estimate the consequences of their interactions for bacterial ecology and evolution, contributing to the recent attention in understanding the multi-party interactions between mobile genetic elements [6, 10, 38–40].

We show that the presence of cells without a CRISPR-Cas system promotes phage survival, consistent with previous research [12]. In addition, we identify the conditions, dependent on both CRISPR-Cas and the phage, that allow segregation loss to increase phage survival. We also describe the driving mechanism for increased phage survival, i.e. the facilitation of phage outbreaks caused by phages that escape CRISPR-Cas. However, most conjugative plasmids encode mechanisms such as partitioning systems or addiction systems, which are effective in keeping segregation loss low [41]. Experiments have shown that plasmids often have a probability of segregation loss of at most 10^*−*3^ or lower [4, 42]. Therefore, our results indicate that plasmids with a CRISPR-Cas system can reliably defend against virulent phages.

Interestingly, CRISPR-Cas systems carried by plasmids generally contain more spacers targeting other plasmids than phages, whereas chromosomal CRISPR-Cas contain more spacers targeting phages [11, 41]. Our results show for CRISPR-Cas systems on plasmids that the relatively low proportion of spacers targeting phages cannot necessarily be explained by a decreased phage protection efficiency. This could support the hypothesis that the bias of CRISPR-Cas systems on plasmids towards spacers targeting other plasmids results from their involvement in plasmid-plasmid competition [11]. Indeed, a bacterial cell can usually be infected by multiple distinct plasmids and if plasmid interactions are detrimental for two co-infecting plasmids, it results in positive frequency selection, ultimately driving the exclusion of the less frequent plasmid from the bacterial population even if the plasmids are otherwise equally fit [19]. Consequently, even though plasmids can be burdensome for bacterial cells, the selection pressure to defend against other plasmids might be higher for a plasmid than the bacterial cell itself. Moreover, the selection pressure to defend against phages is expected to be high for both bacterial cells and plasmids. However, if the bacterial cell is already efficient in defending against phages, the selection pressure to keep anti-phage spacers on plasmids will be weakened and this could result in an increased frequency of spacers targeting competing plasmids.

As our results show that CRISPR-Cas efficiency is for most of the parameter sets studied here independent of its location, phage protection might potentially be outsourced to plasmids. There are several factors that could favour CRISPR-Cas to be carried by plasmids. First, it has been investigated which evolutionary forces drive a gene to be carried by a plasmid rather than the chromosome. [33, 43] Theory predicts that moderately beneficial genes may indeed reside on plasmids rather than on the chromosome, but very beneficial genes are predicted to be carried on the chromosome [33]. Given that CRISPR-Cas can be essential for bacterial survival during a phage attack, it is expected to be located on the chromosome. However, bacteria typically carry several defense mechanisms against phages [3] and some of those defence mechanisms can work synergistically [44, 45]. Consequently, CRISPR-Cas may not always be essential but moderately beneficial for bacterial survival and therefore it could be located on a plasmid. Second, phages that depend on plasmids for infection are more abundant than previously thought [39]. These phages can only infect bacterial cells hosting a plasmid. Consequently, plasmid carriage can confer additional evolutionary pressure on bacteria besides the commonly discussed plasmid fitness effect on bacterial replication. To lower this burden, plasmids may potentially carry CRISPR-Cas to defend against those plasmid dependent phages and ensure plasmid maintenance. Bioinformatic analysis could be used to test whether the phage spacer content of CRISPR-Cas carried by plasmids shows a bias towards plasmid dependent phages. Third, CRISPR-Cas may be favoured to be carried by multi-copy plasmids, in case multiple copies of CRISPR-systems would have synergistic effects and therewith increase phage protection efficiency. Higher phage protection efficiency could be achieved through a higher likelihood for a cell to carry several spacers targeting a given phage (within cell spacer diversity), which is known to be more efficient in blocking phage evolutionary emergence than population-wide spacer diversity [46]. However, it is generally unclear if carrying multiple copies of CRISPR-Cas increases within-cell spacer diversity or confers other benefits.

In summary, we show that CRISPR-Cas on plasmids provides robust protection against virulent phages, showing that plasmids carrying CRISPR-Cas could play a role in phage defence. This highlights the importance of considering the interaction between mobile genetic elements (including their arsenal of immune defences) to understand the ecology and evolution of bacteria in complex environments.

## Data availability

Data and code are available at OSF: https://osf.io/hcvsr/?view_only=553bb579a1e04d7d855b756d22d9dc7e.

## Acknowledgments

We thank Sonja Lehtinen, Claudia Igler and Martin Guillemet for their helpful comments on the manuscript. This project was supported by an ETH Zurich Postdoctoral Fellowship (18-2-FEL-51) received by HC.

## References

1. Vogwill T, MacLean RC. The genetic basis of the fitness costs of antimicrobial resistance: a meta-analysis approach. Evolutionary Applications. 2015;8(3):284–95.

2. Calendar R. The bacteriophages. vol. 2. Oxford University Press; 2006.

3. Georjon H, Bernheim A. The highly diverse antiphage defence systems of bacteria. Nature Reviews Microbiology. 2023:1–15.

4. Lau BT, Malkus P, Paulsson J. New quantitative methods for measuring plasmid loss rates reveal unexpected stability. Plasmid. 2013;70(3):353–61.

5. Rocha EP, Bikard D. Microbial defenses against mobile genetic elements and viruses: who defends whom from what? PLoS biology. 2022;20(1):e3001514.

6. Harrison E, Wood AJ, Dytham C, Pitchford JW, Truman J, Spiers A, et al. Bacteriophages limit the existence conditions for conjugative plasmids. MBio. 2015;6(3):10–1128.

7. Pecota DC, Wood TK. Exclusion of T4 phage by the hok/sok killer locus from plasmid R1. Journal of bacteriology. 1996;178(7):2044–50.

8. Fineran PC, Blower TR, Foulds IJ, Humphreys DP, Lilley KS, Salmond GP. The phage abortive infection system, ToxIN, functions as a protein–RNA toxin–antitoxin pair. Proceedings of the National Academy of Sciences. 2009;106(3):894–9.

9. Dy RL, Przybilski R, Semeijn K, Salmond GP, Fineran PC. A widespread bacteriophage abortive infection system functions through a Type IV toxin–antitoxin mechanism. Nucleic acids research. 2014;42(7):4590–605.

10. Bleriot I, Blasco L, Pacios O, Fernández-Garcia L, Ambroa A, López M, et al. The role of PemIK (PemK/PemI) type II TA system from Klebsiella pneumoniae clinical strains in lytic phage infection. Scientific reports. 2022;12(1):4488.

11. Pinilla-Redondo R, Russel J, Mayo-Muñoz D, Shah SA, Garrett RA, Nesme J, et al. CRISPR-Cas systems are widespread accessory elements across bacterial and archaeal plasmids. Nucleic acids research. 2022;50(8):4315–28.

12. Weissman JL, Holmes R, Barrangou R, Moineau S, Fagan WF, Levin B, et al. Immune loss as a driver of coexistence during host-phage coevolution. The ISME journal. 2018;12(2):585–97.

13. Deveau H, Barrangou R, Garneau JE, Labonté J, Fremaux C, Boyaval P, et al. Phage response to CRISPR-encoded resistance in Streptococcus thermophilus. Journal of bacteriology. 2008;190(4):1390–400.

14. Chabas H, Lion S, Nicot A, Meaden S, van Houte S, Moineau S, et al. Evolutionary emergence of infectious diseases in heterogeneous host populations. PLoS biology. 2018;16(9):e2006738.

15. van Houte S, Ekroth AK, Broniewski JM, Chabas H, Ashby B, Bondy-Denomy J, et al. The diversity-generating benefits of a prokaryotic adaptive immune system. Nature. 2016;532(7599):385–8.

16. Common J, Morley D, Westra ER, van Houte S. CRISPR-Cas immunity leads to a coevolutionary arms race between Streptococcus thermophilus and lytic phage. Philosophical Transactions of the Royal Society B. 2019;374(1772):20180098.

17. Guillemet M, Chabas H, Nicot A, Gatchich F, Ortega-Abboud E, Buus C, et al. Competition and coevolution drive the evolution and the diversification of CRISPR immunity. Nature Ecology & Evolution. 2022:1–9.

18. Chabas H, Müller V, Bonhoeffer S, Regoes RR. Epidemiological and evolutionary consequences of different types of CRISPR-Cas systems. PLoS computational biology. 2022;18(7):e1010329.

19. Igler C, Huisman JS, Siedentop B, Bonhoeffer S, Lehtinen S. Plasmid co-infection: linking biological mechanisms to ecological and evolutionary dynamics. Philosophical Transactions of the Royal Society B. 2022;377(1842):20200478.

20. Makarova KS, Wolf YI, Alkhnbashi OS, Costa F, Shah SA, Saunders SJ, et al. An updated evolutionary classification of CRISPR–Cas systems. Nature Reviews Microbiology. 2015;13(11):722–36.

21. Paez-Espino D, Morovic W, Sun CL, Thomas BC, Ueda Ki, Stahl B, et al. Strong bias in the bacterial CRISPR elements that confer immunity to phage. Nature communications. 2013;4(1):1–7.

22. Heler R, Wright AV, Vucelja M, Doudna JA, Marraffini LA. Spacer acquisition rates determine the immunological diversity of the type II CRISPR-Cas immune response. Cell host & microbe. 2019;25(2):242–9.

23. Garcillán-Barcia MP, de la Cruz F. Why is entry exclusion an essential feature of conjugative plasmids? Plasmid. 2008;60(1):1–18.

24. Chabas H, Nicot A, Meaden S, Westra ER, Tremblay DM, Pradier L, et al. Variability in the durability of CRISPR-Cas immunity. Philosophical Transactions of the Royal Society B. 2019;374(1772):20180097.

25. Garcillán-Barcia MP, de la Cruz F. Why is entry exclusion an essential feature of conjugative plasmids? Plasmid. 2008;60(1):1–18.

26. Johnson P. adaptivetau: Tau-Leaping Stochastic Simulation; 2019. R package version 2.2-3. Available from: https://CRAN.R-project.org/package=adaptivetau.

27. Cao Y, Gillespie DT, Petzold LR. Adaptive explicit-implicit tau-leaping method with automatic tau selection. The Journal of chemical physics. 2007;126(22):224101.

28. Magadán AH, Dupuis ME’, Villion M, Moineau S. Cleavage of phage DNA by the Streptococcus thermophilus CRISPR3-Cas system. PloS one. 2012;7(7):e40913.

29. Keogh BP. Adsorption, latent period and burst size of phages of some strains of lactic streptococci. Journal of Dairy Research. 1973;40(3):303–9.

30. Parada V, Herndl GJ, Weinbauer MG. Viral burst size of heterotrophic prokaryotes in aquatic systems. Journal of the Marine Biological Association of the United Kingdom. 2006;86(3):613–21.

31. Yang Y, Ma R, Yu C, Ye J, Chen X, Wang L, et al. A novel alteromonas phage lineage with a broad host range and small burst size. Microbiology Spectrum. 2022;10(4):e01499–22.

32. Sheppard RJ, Beddis AE, Barraclough TG. The role of hosts, plasmids and environment in determining plasmid transfer rates: A meta-analysis. Plasmid. 2020;108:102489.

33. Lehtinen S, Huisman JS, Bonhoeffer S. Evolutionary mechanisms that determine which bacterial genes are carried on plasmids. Evolution Letters. 2021 06;5(3):290–301.

34. Chen T, Guestrin C. XGBoost: A Scalable Tree Boosting System. In: Proceedings of the 22nd ACM SIGKDD International Conference on Knowledge Discovery and Data Mining. KDD’16. New York, NY, USA: ACM; 2016. p. 785–94.

35. XGBoost Developers. XGBoost Documentation; 2023. Accessed on October 13th, 2023. Available from: https://xgboost.readthedocs.io/en/latest/.

36. Heler R, Wright AV, Vucelja M, Bikard D, Doudna JA, Marraffini LA. Mutations in Cas9 enhance the rate of acquisition of viral spacer sequences during the CRISPR-Cas immune response. Molecular cell. 2017;65(1):168–75.

37. Bradde S, Vucelja M, Tesileanu T, Balasubramanian V. Dynamics of adaptive immunity against phage in bacterial populations. PLoS computational biology. 2017;13(4):e1005486.

38. Castledine M, Newbury A, Lewis R, Hacker C, Meaden S, Buckling A. Antagonistic Mobile Genetic Elements Can Counteract Each Other’s Effects on Microbial Community Composition. Mbio. 2023;14(2):e00460–23.

39. Quinones-Olvera N, Owen SV, McCully LM, Marin MG, Rand EA, Fan AC, et al. Diverse and abundant viruses exploit conjugative plasmids. bioRxiv. 2023:2023-03.

40. Igler C, Schwyter L, Gehrig D, Wendling CC. Conjugative plasmid transfer is limited by prophages but can be overcome by high conjugation rates. Philosophical Transactions of the Royal Society B. 2022;377(1842):20200470.

41. Rodriguez-Beltrán J, DelaFuente J, Leon-Sampedro R, MacLean RC, San Millan A. Beyond horizontal gene transfer: the role of plasmids in bacterial evolution. Nature Reviews Microbiology. 2021;19(6):347–59.

42. Loftie-Eaton W, Bashford K, Quinn H, Dong K, Millstein J, Hunter S, et al. Compensatory mutations improve general permissiveness to antibiotic resistance plasmids. Nature ecology & evolution. 2017;1(9):1354–63.

43. Svara F, Rankin DJ. The evolution of plasmid-carried antibiotic resistance. BMC evolutionary biology. 2011;11:1–10.

44. Dupuis ME, Villion M, Magadán AH, Moineau S. CRISPR-Cas and restriction–modification systems are compatible and increase phage resistance. Nature communications. 2013;4(1):2087.

45. Maestri A, Pursey E, Chong C, Pons BJ, Gandon S, Custodio R, et al. Bacterial defences interact synergistically by disrupting phage cooperation. bioRxiv. 2023:2023–03.

46. Gandon S, Guillemet M, Gatchitch F, Nicot A, Renaud AC, Tremblay DM, et al. Building pyramids against the evolutionary emergence of pathogens. bioRxiv. 2022.

