## Supplement file for "My host’s enemy is my enemy: plasmids carrying CRISPR-Cas as a defence against phages"

Supplementary materials for *My host's enemy is my enemy: plasmids carrying CRISPR-Cas as a defence against phages*

### 1 XGBoost model training and results

We train a statistical model using XGBoost, which can predict, whether the CRISPR-Cas protection efficiency on the plasmid is better, same or worse than the CRISPR-Cas efficiency on the chromosome. The XGBoost classifier is a ensemble tree model constructed from multiple trees. For training the model, we use 70% of our simulation outputs. As our classes "better", "same", "worse" occur with very different frequency, we use weighting of our input data for training our model. Therewith, we try to avoid that the model only learns to predict the most frequent class "same" very well and is not able to predict the less frequent classes "worse" or "better". We therefore assign in the training process more importance in predicting less frequent classes, by weighing data points from each class inversely to their class frequency. We perform hyper parameter-tuning to improve the performance of our model. After we trained our model we test the performance of our model with the remaining 30% of our simulations. The confusion matrix (table 1) shows how often our trained model predicted the classes right for the test data set.

| | Actual classification | | |
|---|---|---|---|
| Predicted classification | better | same | worse |
| better | 438 | 2714 | 97 |
| same | 59 | 14112 | 447 |
| worse | 19 | 2218 | 2655 |

**Table 1.** Confusion matrix showing the predicted classification of the test set using the trained model and the actual classification.

### 2 Spacer diversity hypothesis

The generated population-based spacer diversity is key in successfully defending against a virulent phage outbreaks. We hypothesised that in case CRISPR-Cas is located on a plasmid, segregation loss could decrease the generated population wide spacer diversity and therefore, the chance of bacterial survival. Segregation loss results in the loss of CRISPR-Cas systems with the opportunity to acquire a spacer, which could lead to a lower spacer diversity. To test this, we investigated the bacterial and phage composition at the time where no naive CRISPR-Cas system remains in the population along the lines of Chabas et al., 2022 [1]. After this time point, no new distinct spacer can be acquired in the bacterial population and the frequency of existing spacers only changes via competition and growth dynamics.

First of all, we test if the probability to acquire at least one spacer is altered by segregation loss and we observe no trend between segregation loss and the probability to acquire at least one spacer (figure S1A). As the probability to acquire at least on spacer is comparable, we investigate in the following only simulations where at least one spacer was acquired. We want to investigate the spacer diversity at the time point where no naive CRISPR-Cas system remains in the population, hereafter called initial spacer diversity. We use two measures to quantify the initial spacer diversity: the richness, and the inverse Simpson index. The richness quantifies how many distinct spacers are acquired. The inverse Simpson's index is a measure of diversity, which also takes into account the frequency of each spacer in the population. The inverse Simpson's Index $\frac{1}{D}$ is defined as follows:

$$\frac{1}{D} = \frac{1}{\sum_{i=1}^{n_s} p_i^2} \tag{1}$$

where $p_i$ is the bacterial genotype $i$, i.e. bacteria with a CRISPR-Cas system carrying phage spacer $i$. The inverse Simpson's index takes on values close to 1 if the bacterial population is dominated by a single spacer and therefore shows little diversity. The higher the inverse Simpson's index the more equal is the frequency of different occurring bacterial genotypes and therefore the population composition is more diverse. We observe that the initial spacer diversity is not influenced by segregation loss for both measures of diversity (figure S1C,D). To exclude effects of a different number of phage virons at initial spacer diversity, we verified that the phage population has a similar population size (figure S1B). We therefore conclude that a decreased generated population-based spacer diversity cannot explain why segregation loss leads to a decreased CRISPR-Cas protection efficiency.

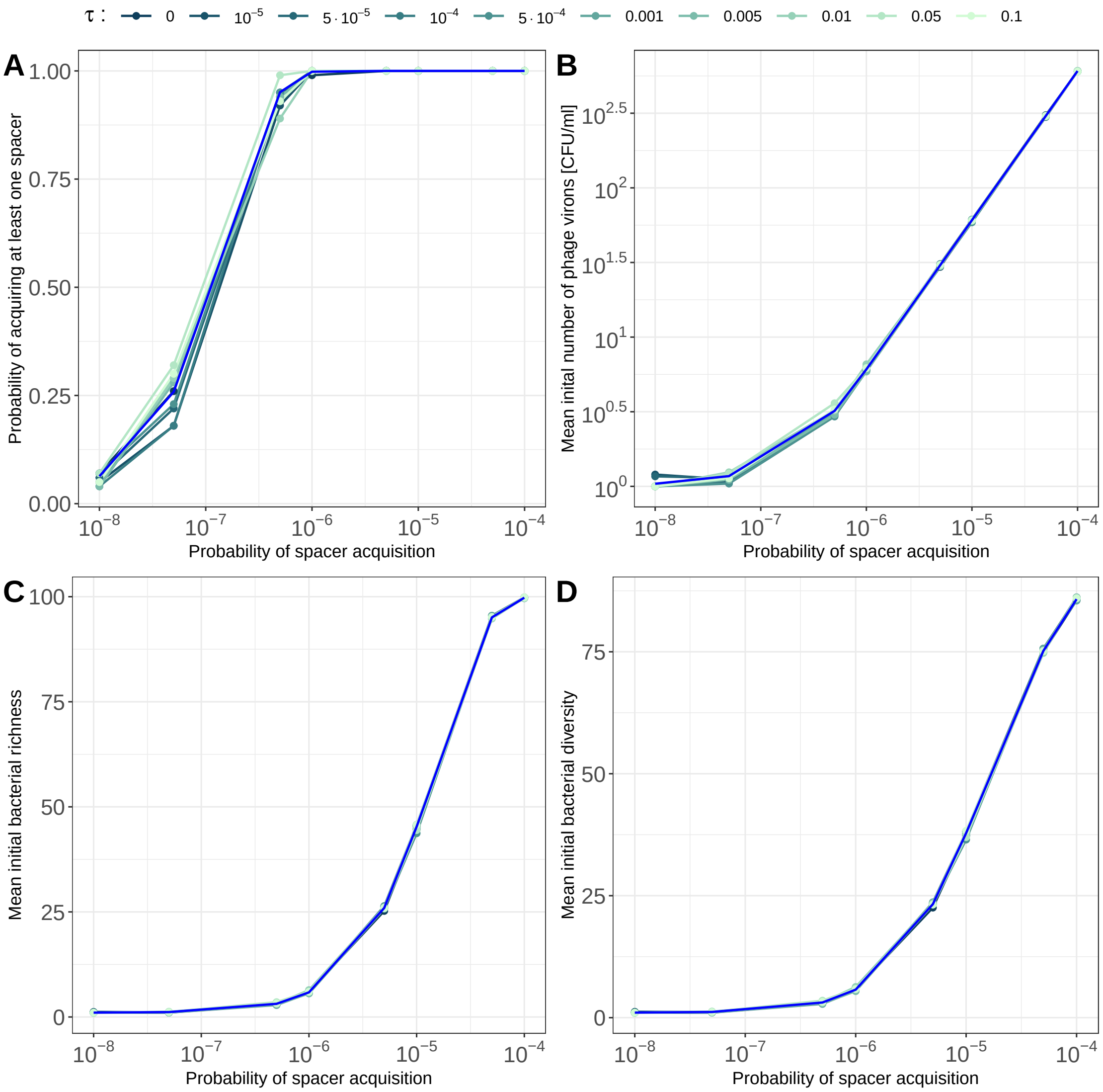


**Figure S1. Segregation loss has no impact on initial spacer diversity.** Simulations for a plasmid with a cost $c = 0.05$, a conjugation rate $\gamma = 10^{-14}$ and varying probabilities of segregation loss $\tau$ are shown in shades of mint. Results for the chromosome are indicated with a blue line. Results are shown for different probabilities of spacer acquisition $\alpha$. **A** Probability of acquiring at least one spacer before the susceptible bacterial population dies out. **B** Mean initial number of phage virons at initial spacer diversity for cases where at least one spacer was acquired. **C** Mean initial bacterial richness at initial spacer diversity for cases where at least one spacer was acquired. **D** Mean initial bacterial diversity, calculated via the inverse Simpson's index (equation 1), for cases where at least on spacer was acquired.

### 3 Phage evolution hypothesis

To test whether segregation loss favours outbreaks of phage mutants, we investigate if segregation loss increases the proportion of phage mutants after a phage outbreak. To do so we plot, for all simulations in which the phage survives, the mean proportion of phages with mutated protospacers at the beginning of the phage outbreak and at the end of the phage outbreak. For plotting the proportion of escape phages at the beginning of the outbreak, we choose the time point at which initial spacer diversity is reached, i.e. no naive CRISPR-Cas system remains in the population. We observe that at the beginning of the phage outbreak the proportion of phage mutants is very small, regardless of the extent of segregation loss (Supplementary Figure S2A). However, at the end of the phage outbreak, we observe that with high segregation loss, the phage population is dominated by phages with mutated protospacers (Supplementary Figure S2 B). This confirms that segregation loss favours outbreaks of phage mutants (*phage evolution hypothesis*).

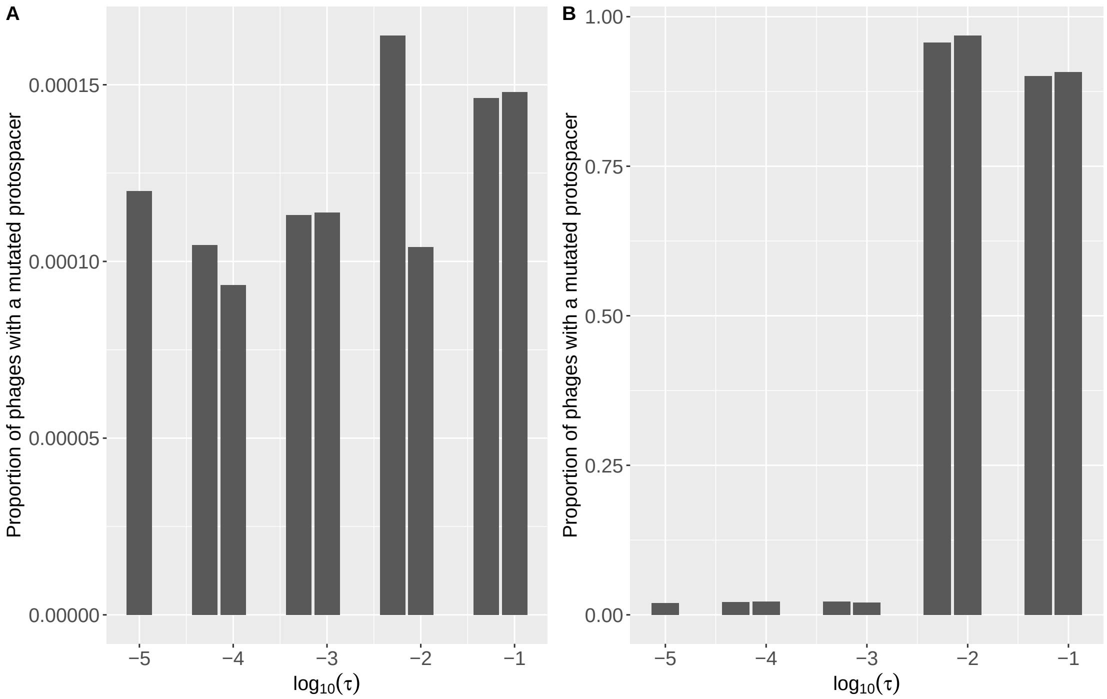


**Figure S2. Segregation loss favours phage outbreaks of escape mutants.** Impact of segregation loss on the mean proportion of escape phages once initial spacer diversity is reached (**A**) and at the end of the phage outbreak (**B**) for cases where the phage survived. The conjugation rate, plasmid fitness effect, phage protospacer mutation probability and phage burst sizes are fixed at $\gamma = 10^{-14}$, $c = 0.05$, $\mu = 3.4 \cdot 10^{-7}$, $b = 190$ and $b_m = 179$.

### References

1. Chabas H, Müller V, Bonhoeffer S, Regoes RR. Epidemiological and evolutionary consequences of different types of CRISPR-Cas systems. PLoS computational biology. 2022;18(7):e1010329.
